# Lineages derived from *Cryptococcus neoformans* type strain H99 support a link between the capacity to be pleomorphic and virulence

**DOI:** 10.1101/2022.02.09.479843

**Authors:** Kenya E. Fernandes, James A. Fraser, Dee A. Carter

**Affiliations:** School of Life and Environmental Sciences, University of Sydney, Sydney, NSW, Australia; The University of Queensland, School of Chemistry and Molecular Biosciences, Australian Infectious Diseases Research Centre, St Lucia, QLD, Australia; Sydney Institute for Infectious Diseases, University of Sydney, Sydney, NSW, Australia

## Abstract

The pathogenic yeast *Cryptococcus neoformans* causes nearly 200,000 deaths annually in immunocompromised individuals. *Cryptococcus* cells can undergo substantial morphological change during mammalian infection, including increased capsule and cell size, the release of shed capsule, and the production of titan (> 10 μm), micro (< 2 μm) and irregular cells. We examined phenotypic variation under conditions designed to simulate *in vivo* stress in a collection of nine lineages derived from the *C. neoformans* type strain H99. These lineages are highly genetically similar but have a range of virulence levels. Strains from hypervirulent lineages had a larger average capsule size, greater variation in cell size, and an increased production of micro cells and shed capsule. We tested whether disruption of *SGF29*, which encodes a component of the SAGA histone acetylation complex that has previously been implicated in the hypervirulence of some lineages, might also have a role in the production of morphological variants. Deletion of *SGF29* in a lineage with intermediate virulence substantially increased its production of micro cells and released capsule, consistent with a switch to hypervirulence. We further examined *SGF29* in a set of 52 clinical isolates and found loss-of-function mutations were significantly correlated with patient death. Expansion of a TA repeat in the second intron of *SGF29* was positively correlated with cell and capsule size, suggesting it may also affect Sgf29 function. This study extends the evidence for a link between pleomorphism and virulence in *Cryptococcus*, with a likely role for epigenetic mechanisms mediated by SAGA-induced histone acetylation.

**IMPORTANCE:** Cryptococcosis is a devastating cause of death and disease worldwide. During infection, *Cryptococcus* cells can undergo substantial changes to their size and shape. In this study, we used a collection *C. neoformans* strains that are highly genetically similar but possess differing levels of virulence to investigate how morphological variation aligns with virulence. We found hypervirulent strains on average had larger capsules and greater variation in cell size, and also produced more micro cells and shed capsule. These hypervirulent strains possessed a mutation in *SGF29*, which encodes a component of the SAGA complex involved in epigenetic regulation. Analysis of the *SGF29* gene in a set of clinical isolates found strains with loss-of-function mutations were associated with higher patient death rates. The capacity to vary appears to be linked with virulence in *Cryptococcus*, and this can occur in the absence of genetic variation via epigenetic mechanisms.

## INTRODUCTION

*Cryptococcus neoformans* is an encapsulated yeast pathogen that primarily infects immunocompromised people and can cause severe meningoencephalitis (1). Despite progress in reducing the incidence and burden of disease through improved diagnostic strategies and access to antiretroviral therapy, the mortality rate associated with cryptococcosis still remains unacceptably high (2). In addition to a paucity of effective and widely available antifungal drugs (3), there is incomplete knowledge of the interplay between pathogen and host with which to inform clinical management (1). Cryptococcosis can be associated with a range of clinical presentations and outcomes, and this is determined in part by variations in the infecting strain (3). During mammalian infection, genetically identical Cryptococcus cells can undergo substantial phenotypic variation including the production of titan, micro, and irregular cell variants, a dramatic increase in capsule size, and the shedding of capsule (4). These changes to the size and shape of cells are frequently observed in clinical samples, however relatively little is known about the full variety of morphologies that cryptococcal cells can produce and how this affects their pathogenicity.

Previously, we investigated phenotypic variation in a collection of 70 isolates from HIV/AIDS patients from Botswana with cryptococcosis and found significant correlations between phenotypic and clinical data (5). In particular, the combined presence of giant cells (which we defined as cells that are in the size range of titan cells [> 15 μm] but cannot be definitively described as titan cells as polyploidy and increased cell wall chitin (6, 7) have not been determined), micro cells (≤ 1 μm) and shed capsule, while rare, was positively correlated with patient death, indicating that the capacity for variation might play a role in virulence. While interesting correlations were uncovered by this study, it was limited to being mostly descriptive, with the high level of diversity present in both the cryptococcal isolates and host populations confounding efforts to tease apart cause and effect.

*Cryptococcus neoformans* strain H99 was first isolated in 1978 and has since become the most widely used C. neoformans reference strain globally. During storage, passage, and sub-culture in various labs around the world, distinct H99 lineages have arisen with varying levels of virulence (Fig. 1A) (8). Complete sequencing of strains representing these lineages has revealed a small number of unique mutations among them. Notably, the hypervirulent H99L lineage strains have a 734 bp deletion in the gene encoding Sgf29, a component of the SAGA histone acetylation complex. This complex is involved in controlling gene transcription through remodelling chromatin structure and its loss has been shown to lead to a reduction in histone H3K9 acetylation across the genome (9). In H99L and its derivatives, the disruption of Sgf29 has been implicated in the switch to the hypervirulent phenotype and Sgf29 loss-of-function mutations have been identified in clinical isolates (9). Additionally, a group of less virulent H99 lineages possess a mutation in *LMP1* that compromises fecundity and virulence (8).

**Figure 1:**
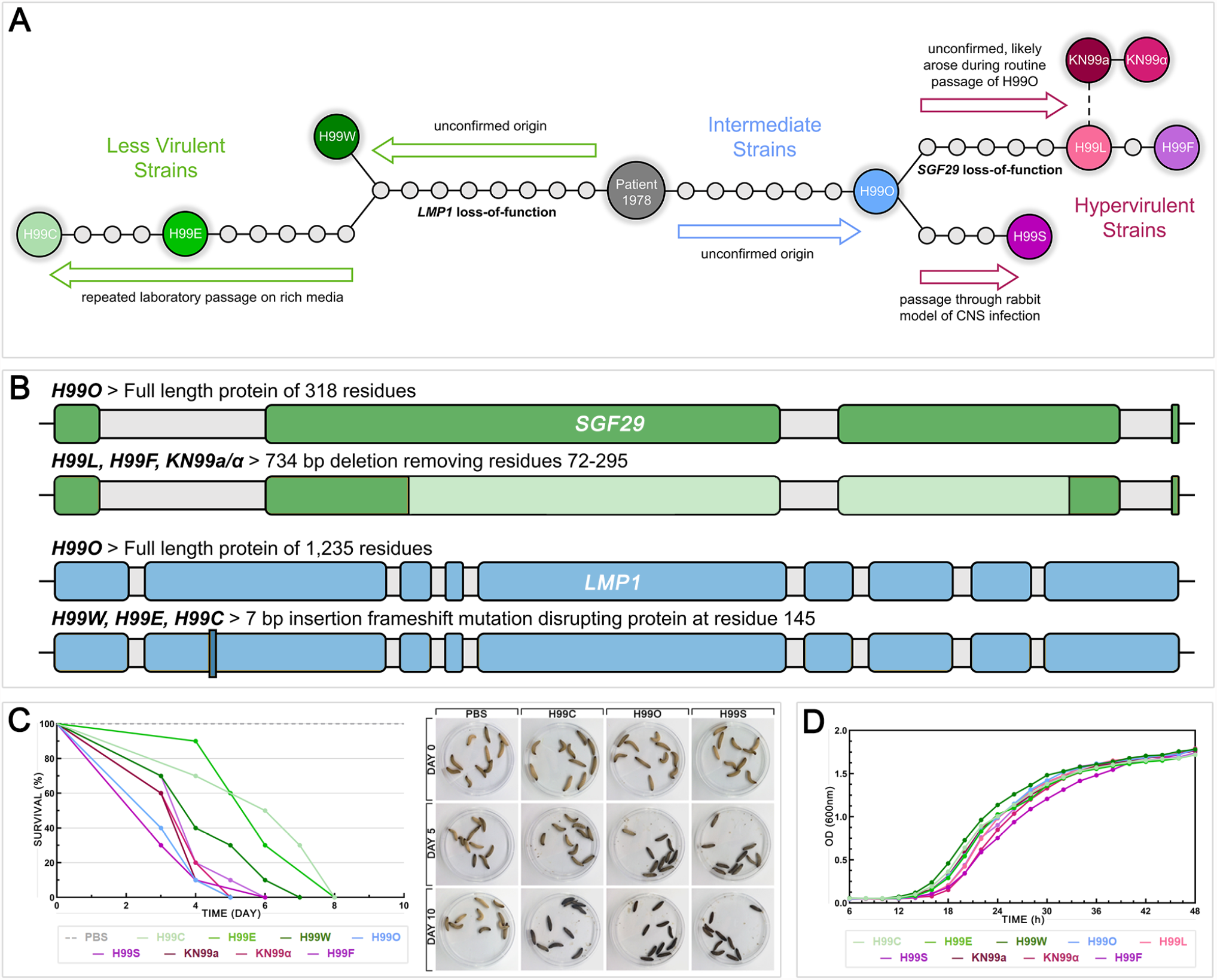
Origins of the *C. neoformans* H99 strains used in this study and their comparative virulence and growth dynamics. **(A)** The relationship between the H99 derivative strains used in this study. Mutations separating strains are shown by circles and known or hypothesised directions of evolution are shown by arrows. Figure adapted from Arras et al. 2017. **(B)** Diagram of the *SGF29* and *LMP1* genes showing disruptions found in some H99 derivative strains. Lighter coloured sections indicate areas of deletion, darker coloured sections indicate areas of insertion. Grey shading indicates introns. **(C)** Survival of *G. mellonella* larvae infected with an inoculum of 10^8^ cells of each H99 strain or mock-infected with PBS over 10 days; n=10 for each treatment group. Left panel shows a representative survival plot (one of two independent replicates that gave similar results); right panel shows photographs of representative larvae at days 0, 5, and 10 that had been mock-infected (PBS) or infected with a less virulent (H99C), intermediate (H99O), or hypervirulent (H99S) strain. Treatment groups were compared by log-rank test; H99C (p ≤0.017) and H99E (p ≤0.019) survived significantly longer than H99O, H99S, H99L, KN99**a**, KN99α, and H99F. **(D)** Growth curves of H99 strains incubated at 30 °C with 180 rpm shaking averaged across three independent replicates, showing no substantial differences between strains.

This set of very closely related H99 strains comprises a valuable resource for determining a link between virulence and phenotype in a background of high genetic similarity. Here, we examined variation in capsule and cell size in this strain set under in vitro conditions designed to simulate in vivo stress and provide evidence that the capacity for phenotypic variation is associated with increased virulence in these strains. We determined growth characteristics and antifungal susceptibility and confirmed that, in most respects, the strains were very similar although hypervirulent strains showed slightly increased susceptibility to some drugs. Finally, we investigated the impact of mutations in *SGF29* on strains in the H99 derivative set and in the set of clinical isolates from Botswana that was investigated in our prior work.

## RESULTS

### H99 derivative strains display expected virulence and growth dynamics

The relationship between the nine H99 lineage strains used in this study is shown in Fig. 1A-B. H99O is believed to be one of the closest remaining relatives of the original H99 strain and is considered to have intermediate virulence. Hypervirulent strains H99L and H99F possess the 734 bp deletion in the gene Sgf29 that appears to play a role in conferring the hypervirulent phenotype (9). The remaining hypervirulent strain H99S does not possess this mutation and is theorised to have acquired hypervirulence in an independent manner (9). H99W, H99E, and H99C comprise the less virulent strains, and possess the mutation in *LMP1* (8). Two additional closely related strains are included in the set: the congenic mating type pair KN99**a** and KN99α. These strains were created through repeat backcrossing between H99F and an unrelated MAT**a** mating type isolate (10) and also display hypervirulence.

On receipt of the strains, we tested their virulence in *Galleria mellonella* (wax moth) larvae to confirm that they were behaving as expected (Fig. 1C). Larvae infected with less virulent strains H99C, H99E and H99W survived 1-3 days longer than larvae infected with intermediate or hypervirulent strains, with H99C (p ≤0.017) and H99E (p ≤0.019) reaching significance against H99O, H99S, H99L, KN99**a**, KN99α, and H99F. These results were consistent with those previously obtained using the *G. mellonella* model (8). Analysis of the growth dynamics of strains under standard growth conditions (SDB at 30 °C) over 48 hours found no substantial difference among them, indicating that all strains were healthy and capable of similar, normal growth (Fig. 1D). Antifungal susceptibility six commonly used antifungal drugs (amphotericin B [AMB], nystatin [NYS], fluconazole [FLC], itraconazole [ITC], voriconazole [VRC], and flucytosine [5FC]) found all strains were equally susceptible to polyene drugs AMB and NYS, while the hypervirulent strains H99S, KN99**a**, KN99α and H99F had increased susceptibility to azole drugs FLC, ITC and VRC, and less virulent strains were 2-fold less susceptible to 5FC (Table 1).

**Table 1:** Strains used in this study with details of cell and capsule size, production of variant phenotypes and antifungal susceptibilities ^a^.

| Strain | Cell Diameter (μm) |  | Capsule Thickness (μm) |  | Irregular Cells (%) | Giant Cells <sup>b</sup> | Micro Cells <sup>b</sup> | Released Capsule <sup>b</sup> | Clustered Capsule <sup>b</sup> | Minimum Inhibitory Conc. (μg/ml) <sup>c</sup> |  |  |  |  |  |
| --- | --- | --- | --- | --- | --- | --- | --- | --- | --- | --- | --- | --- | --- | --- | --- |
|  | Mean | Std Dev | Mean | Std Dev |  |  |  |  |  | AMB | NYS | FLC | ITC | VRC | 5FC |
| Less Virulent |  |  |  |  |  |  |  |  |  |  |  |  |  |  |  |
| H99C | 5.31 | 0.79 | 2.19 | 0.70 | <5 | – | – | – | + | 0.5 | 4 | 4 | 0.25 | 0.125 | 8 |
| H99E | 5.64 | 0.84 | 2.27 | 0.73 | <5 | + | – | + | – | 0.5 | 4 | 4 | 0.25 | 0.125 | 8 |
| H99W | 5.75 | 0.75 | 1.71 | 0.68 | <5 | – | – | + | + | 0.5 | 4 | 4 | 0.25 | 0.125 | 8 |
| All Less Virulent Strains | Mean | Var | Mean | Var | Strains with Phenotype (%) |  |  |  |  | Geomean |  |  |  |  |  |
|  | 5.57 | 0.03 | 2.06 | 0.06 | 100 | 33 | 0 | 67 | 67 | 0.5 | 4 | 4 | 0.25 | 0.125 | 8 |
| Intermediate Virulence |  |  |  |  |  |  |  |  |  |  |  |  |  |  |  |
| H99O | 5.87 | 1.01 | 3.30 | 0.88 | <5 | + | + | + | + | 0.5 | 4 | 4 | 0.25 | 0.125 | 4 |
| Hypervirulent |  |  |  |  |  |  |  |  |  |  |  |  |  |  |  |
| H99S | 5.86 | 1.21 | 3.07 | 0.77 | <5 | – | + | ++ | +++ | 0.5 | 4 | 2 | 0.125 | 0.063 | 4 |
| H99L | 5.65 | 1.37 | 3.33 | 0.76 | <5 | – | + | ++ | +++ | 0.5 | 4 | 4 | 0.25 | 0.125 | 4 |
| KN99a | 4.72 | 1.18 | 3.02 | 0.85 | <10 | – | +++ | +++ | + | 0.5 | 4 | 1 | 0.125 | 0.031 | 4 |
| KN99α | 4.75 | 1.21 | 2.90 | 0.73 | <10 | – | +++ | +++ | ++ | 0.5 | 4 | 1 | 0.125 | 0.063 | 4 |
| H99F | 5.69 | 1.48 | 2.96 | 0.83 | <5 | – | ++ | ++ | +++ | 0.5 | 4 | 4 | 0.125 | 0.125 | 4 |
| All Hypervirulent Strains | Mean | Var | Mean | Var | Strains with Phenotype (%) |  |  |  |  | Geomean |  |  |  |  |  |
|  | 5.33 | 0.24 | 3.06 | 0.02 | 100 | 0 | 100 | 100 | 100 | 0.5 | 4 | 2 | 0.14 | 0.07 | 4 |
| H99O SGF29 Knockouts |  |  |  |  |  |  |  |  |  |  |  |  |  |  |  |
| H99O.2 | 5.41 | 1.34 | 3.31 | 1.13 | <5 | – | + | + | + | 0.5 | 4 | 4 | 0.25 | 0.125 | 4 |
| H99OΔ | 5.53 | 1.32 | 3.23 | 0.60 | <5 | – | ++ | ++ | ++ | 0.5 | 4 | 4 | 0.25 | 0.125 | 4 |
| H99OΔ+ | 5.26 | 1.79 | 3.32 | 1.49 | <5 | – | ++ | ++ | ++ | 0.5 | 4 | 4 | 0.25 | 0.125 | 4 |
| All H99O KO Strains | Mean | Var | Mean | Var | Strains with Phenotype (%) |  |  |  |  | Geomean |  |  |  |  |  |
|  | 5.40 | 0.01 | 3.29 | 0.00 | 100 | 0 | 100 | 100 | 100 | 0.5 | 4 | 4 | 0.25 | 0.125 | 4 |
| H99S SGF29 Knockouts |  |  |  |  |  |  |  |  |  |  |  |  |  |  |  |
| H99S.2 | 5.28 | 1.14 | 3.35 | 0.63 | <5 | – | + | + | ++ | 0.5 | 4 | 2 | 0.125 | 0.063 | 4 |
| H99SΔ | 5.27 | 1.11 | 3.32 | 0.57 | <5 | – | + | + | ++ | 0.5 | 4 | 4 | 0.25 | 0.125 | 4 |
| H99SΔ+ | 5.56 | 1.16 | 3.14 | 0.69 | <5 | – | + | + | ++ | 0.5 | 4 | 4 | 0.25 | 0.125 | 4 |
| All H99S KO Strains | Mean | Var | Mean | Var | Strains with Phenotype (%) |  |  |  |  | Geomean |  |  |  |  |  |
|  | 5.37 | 0.02 | 3.27 | 0.01 | 100 | 0 | 100 | 100 | 100 | 0.5 | 4 | 3.17 | 0.20 | 0.10 | 4 |

<sup>a</sup> Giant, micro, and irregular cells were excluded from calculations of cell diameter and capsule thickness.
<sup>b</sup> The relative frequency of each morphological variant is shown for individual strains; see Fig. 3A for scoring process.
<sup>c</sup> AMB, amphotericin B; NYS, nystatin; FLC, fluconazole; ITC, itraconazole; VRC, voriconazole; 5FC, 5-fluorocytosine

### Hypervirulent strains produce cells with greater variation in size and larger capsules

Capsule thickness and cell body diameter were measured for 100 individual cells per strain after growth in capsule-inducing conditions (DMEM with CO_2_ at 37 °C for 5 days) (Table 1; Fig. 2A-B). Fig. 2C shows the microscopic morphology of typical cells from each strain. The capsule thickness of less virulent strains was on average approximately 50% smaller than intermediate and hypervirulent strains (p<0.0001). Hypervirulent strain H99L and intermediate strain H99O had the largest average capsules, with H99L reaching significance against all other hypervirulent strains (p ≤0.0165). F test analysis revealed that within the less virulent strains there was considerably less variation in capsule thickness, with H99W and H99C having significantly less variation than intermediate strain H99O (p=0.0145 and p=0.0276, respectively). For average cell body diameter, hypervirulent strains KN99**a** and KN99α were significantly smaller than all other intermediate and hypervirulent strains (p<0.0001) as well as the less virulent strains (p ≤0.0002). Intermediate strain H99O and hypervirulent strain H99S had the largest cell body diameters, while the remaining less virulent and hypervirulent strains ranked in the middle. Once again, there was much less variation in cell diameter in less virulent strains compared to intermediate strain H99O (p ≤0.0028) and all hypervirulent strains (p ≤0.0001). Hypervirulent strains displayed the most cell diameter variation in general, with significance reached against intermediate strain H99O by H99F (p=0.0002) and H99L (p=0.0029).

**Figure 2:**
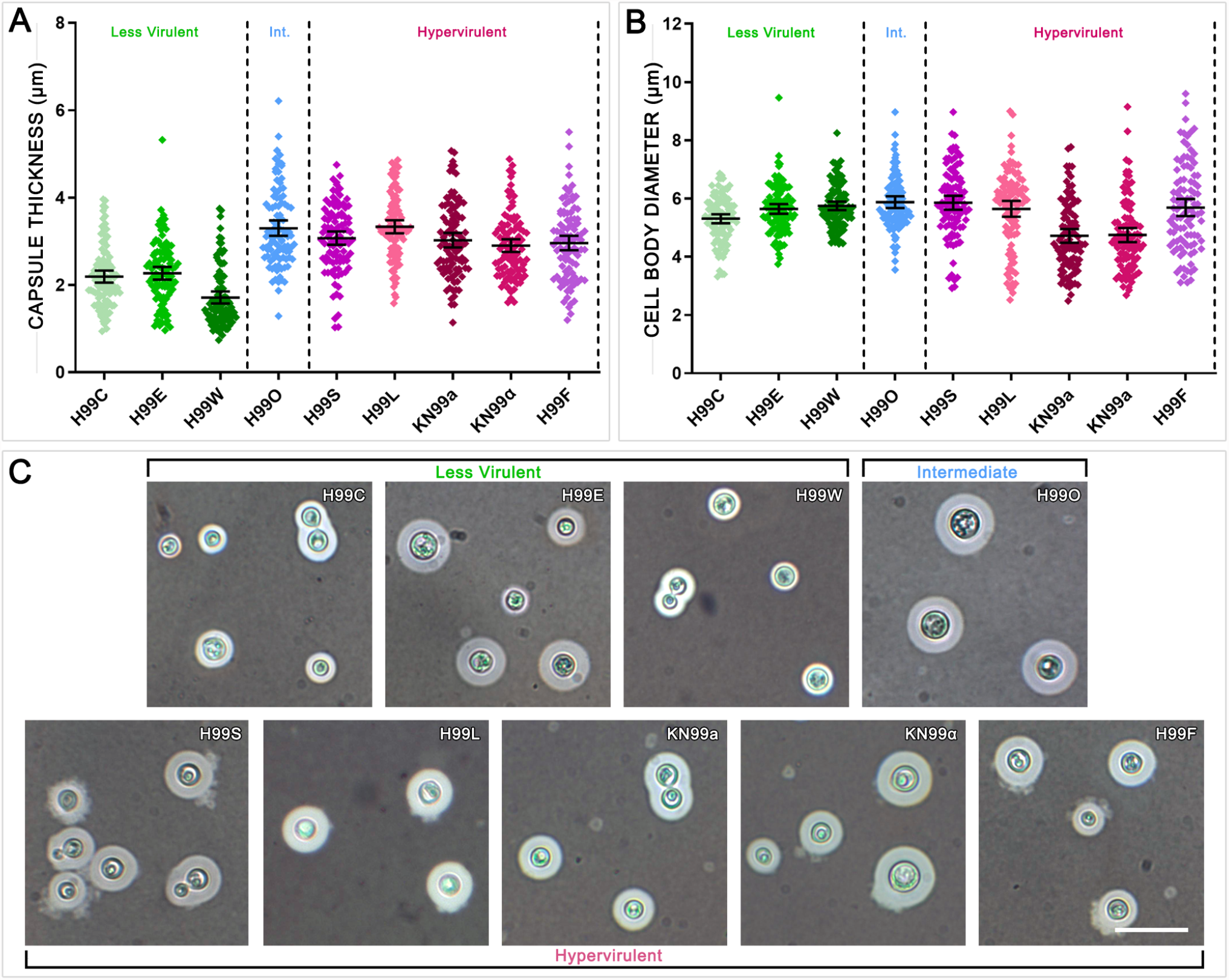
The less virulent H99 strains produce significantly smaller capsules following growth under capsule-inducing conditions. **(A)** Capsule thickness and **(B)** cell body diameter of 100 individual cells of each strain grown in DMEM with 5% CO_2_ at 37°C for 5 days. Strains were compared using two-tailed unpaired t tests with Welch’s correction (significance described in results). Error bars show the mean ± 95% confidence interval. Int. = Intermediate. Giant, micro, and irregular cells were excluded from this analysis. **(C)** Indian ink preparations of each strain showing variation in capsule and cell size. Scale bar = 15 μm.

### Hypervirulent strains produce morphological variants with higher frequency

Reports on the typical size of cryptococcal cells vary, but they are generally considered to be from 4 to 10 μm in diameter (11, 12). Cells substantially larger than average have previously been defined as titan/giant cells, however the exact size definition has varied, including a cell body size of >10 μm, a cell body size >15 μm, and total cell size including capsule >30 μm (13–16). Likewise, cells substantially smaller than average have previously been defined as micro cells, typically defined as a cell body size of < 1 μm (17). The current study found that following capsule induction, the average cell body size of H99 derivative strains ranged from 4.72 μm (KN99α) to 5.87 μm (H99O) with >90% of cells falling between 3 and 8 μm. Based on this, we have defined any cell with a cell body size >10 μm as a giant cell and any cell with cell body size < 2 μm as a micro cell, as these cell sizes were distinctly separate from average cells. The frequency with which these variant cell types were observed in each strain, along with the frequency of extracellular capsule seen either clustered tightly around groups of cells (clustered capsule) or released into the media (released capsule), is shown in Table 1 and Fig. 3. Elongated and irregularly shaped cells were also seen across all strains, but these comprised a small subset (<5%) of the total population of cells, except in KN99**a** and KN99α where they were around twice as common (<10%) (Table 1).

**Figure 3:**
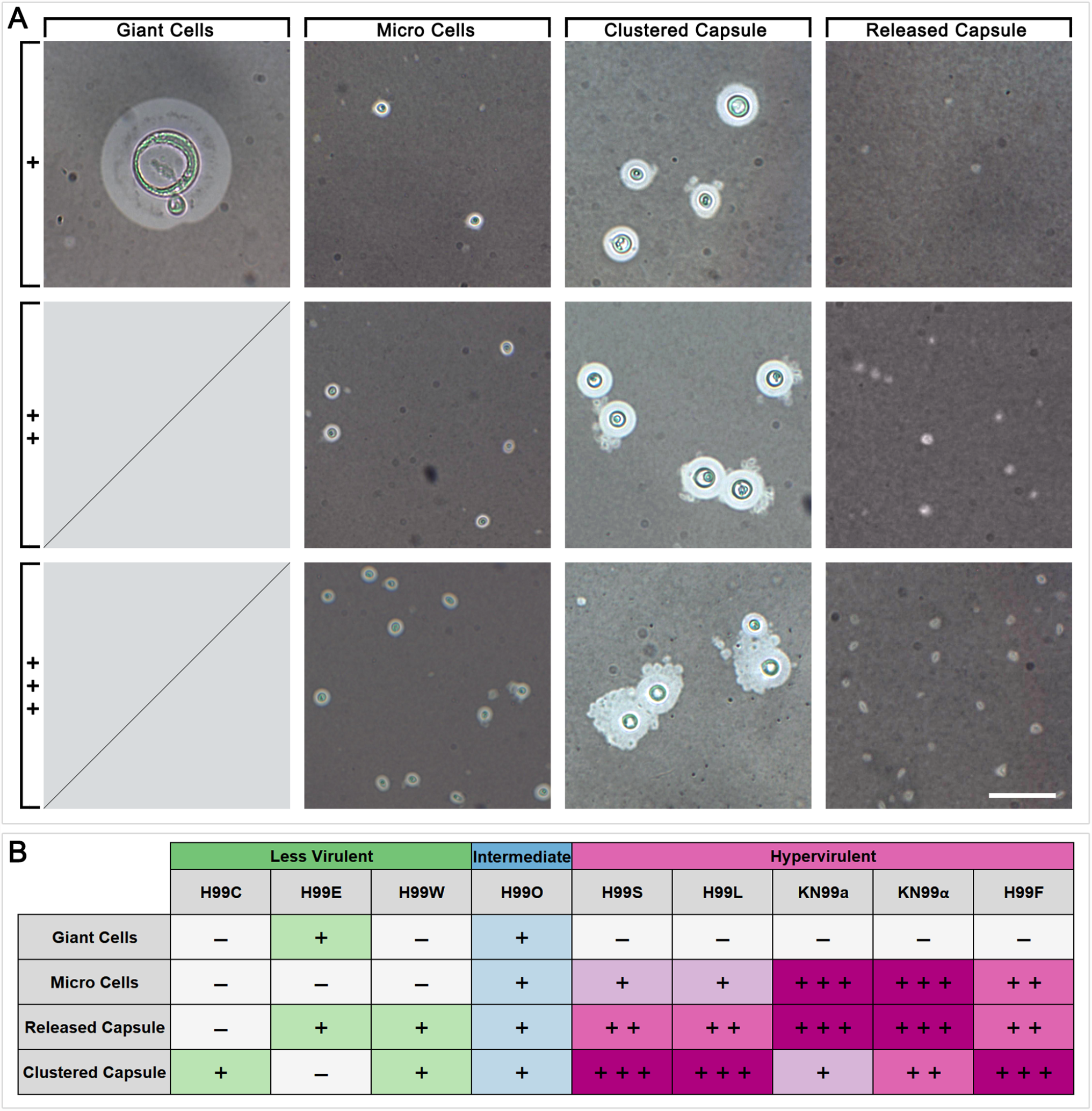
Hypervirulent H99 strains produce morphological variants at a much higher frequency than intermediate and less virulent strains following growth under capsule-inducing conditions. **(A)** Indian ink preparations of H99 strains showing morphological variants produced following growth in DMEM with 5% CO_2_ at 37°C for 5 days, including giant cells larger than 10 μm, micro cells smaller than 2 μm, extracellular capsule clustered around cells (Clustered Capsule), and extracellular capsule released into the media (Released Capsule). Scoring on the left shows the approximate relative frequency of morphological variants seen in a typical field of view, as presented in **(B)**.

Giant cells were observed very infrequently and only in two strains: H99E and H99O. Micro cells were observed in all intermediate and hypervirulent strains and were particularly abundant in KN99**a** and KN99α but were absent from the less virulent strains. Released capsule and clustered capsule were similarly observed in all intermediate and hypervirulent strains. Unlike micro cells, these were each present in two of the three less virulent strains, however they were low in abundance. Clustered capsule was particularly abundant in hypervirulent strains H99S, H99L and H99F, while released capsule was abundant in KN99**a** and KN99α. Overall, each of the hypervirulent strains were observed to produce three out of four of these variant phenotypes and with generally much higher frequency relative to intermediate and less virulent strains. Intermediate strain H99O produced all four variants but with lower relative frequency, while less virulent strains produced between one and two variants with similarly low frequency.

### Electron microscopy reveals defined internal structures in the various cell types

Serial block-face scanning electron microscopy (SBF-SEM) was used to investigate the internal morphology of the giant, micro and irregular cells to confirm that they possess attributes of functional cells. SBF-SEM is a powerful imaging technique that generates serial images through a cell by repeated cutting, raising, and imaging of the sample block, however, the trade-off is that due to electron charging it can have a reduction in resolution compared to transmission electron microscopy (TEM) (18). Supplementary Fig. 1A-D shows representative regular, giant, micro, and irregular cells with slices taken at 10 nm intervals. The giant cell has a thicker cell wall (820 nm) compared to the regular (510 nm) and micro (330 nm) cells, while the cell wall of the irregular cell was particularly thin (260 nm). The giant cells also appeared to have numerous small vesicles. The internal structure of the micro and irregular cells appeared very similar to that of regular cells, with distinguishable vacuoles and other organelles. SEM was used to further visualise the external appearance of micro cells and shed capsule, and clearly demonstrated that these were distinct structures (Supplementary Fig. 1E-G).

### Deletion of *SGF29* increases the production of morphological variants in the H99O lineage but not the H99S lineage

Given that a 734 bp deletion in *SGF29* in the H99L lineage has been implicated in the switch to hypervirulence, we investigated the effect of *SGF29* deletion on the production of cell phenotype variants. To do so, we used *SGF29* knockout and complemented strains in the background of a strain with intermediate virulence (H99O) and a strain with hypervirulence independent of the *SGF29* deletion (H99S) described in (9): H99O.2 and H99S.2, which were used in the creation of the deletion strains and are presumed to be identical copies of H99O and H99S, respectively; *sgf29Δ* ^H99O^ (here named H99OΔ) and *sgf29Δ* ^H99S^ (named H99SΔ), which are *sgf29Δ* deletion mutants created by biolistic transformation; and *sgf29Δ* + *SGF29* ^H99O^ (here named H99OΔ+) and *sgf29Δ* + *SGF29* ^H99S^ (named H99SΔ+), which are *sgf29Δ* deletion mutants complemented with wild-type *SGF29* (9) (Table 1).

Growth curves of wild-type, mutant, and complemented strains under standard growth conditions (SDB at 30 °C) over 48 hours confirmed that there were no significant differences or abnormalities in the growth rate of these strains (Fig. 4A-B). Antifungal susceptibility testing showed that H99SΔ and H99SΔ+ had 2-fold reduced susceptibility to FLC, ITC, and VRC while mutant and complement strains of H99O displayed no differences compared to the wild-type (Table 1). After growth under capsule-inducing conditions in DMEM with CO_2_ at 37 °C for 5 days, there were no significant differences in capsule thickness or cell body diameter between H99O.2 and either H99OΔ or H99OΔ+ and between H99S.2 and either H99SΔ or H99SΔ+ (Fig. 4C-D). H99O.2 and H99S.2 produced the same types and amounts of morphological variants as H99O and H99S, respectively, indicating that these traits are stable characteristics of the strain. In the H99O lineage, the H99OΔ mutant produced substantially increased amounts of micro cells, released capsule, and clustered capsule and this was retained in the complemented H99OΔ+ strain. In contrast, in the H99S lineage, the H99SΔ mutant and its complement strain H99SΔ+ displayed no appreciable difference in the presence or frequency of morphological variants.

**Figure 4:**
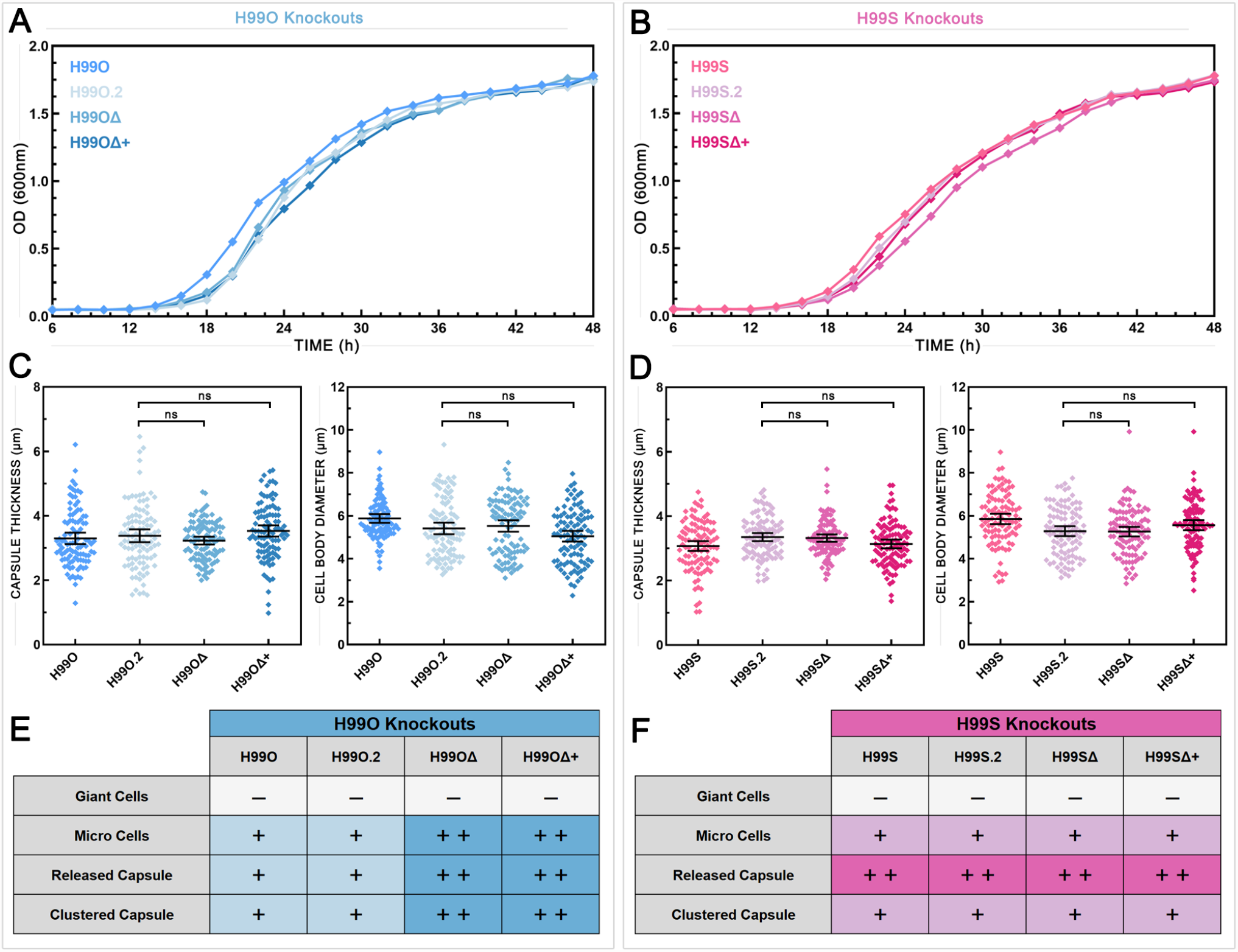
Deletion of *SGF29* increases the production of morphological variants by H99O but not by H99S. Growth curves of wild-type and *sgf29* knockout strains of **(A)** H99O and **(B)** H99S incubated at 30 °C with 180 rpm shaking averaged across three independent replicates showing no substantial differences among strains. Capsule thickness (left panels) and cell body diameter (right panels) of wild-type and *sgf29* knockout strains of **(C)** H99O and **(D)** H99S grown in DMEM with 5% CO_2_ at 37°C for 5 days. Strains were compared using two-tailed unpaired t tests with Welch’s correction. Error bars show the mean ± 95% confidence interval, n = 100. The relative frequency of morphological variants in wild-type and *sgf29* knockout strains of **(E)** H99O and **(F)** H99S; see Fig. 3A for scoring process. H99O.2 and H99S.2 are independently sourced and presumed identical copies of H99O and H99S, respectively. H99OΔ = *sgf29Δ* ^H99O^; H99OΔ+ = *sgf29Δ* + *SGF29* ^H99O^; H99SΔ = *sgf29Δ* ^H99S^; H99SΔ+ = *sgf29Δ* + *SGF29* ^H99S^.

### In clinical isolates, loss-of-function mutations in *SGF29* correlate with patient death

Based on the association between *sgf29Δ*, virulence, and cell phenotypes seen in the H99 strains, we extended our analysis of *SGF29* to a set of clinical isolates from Botswana that had been fully sequenced and extensively analysed for cell morphology and associations with clinical parameters (5). Using whole genome sequencing data (19) (accession numbers in Supplementary Table S1), we examined *SGF29* in 52 *C. neoformans* strains across 4 genotypes (VNI n=16; VNII n=2; VNBI n=25; VNBII n=9). Three strains, NRHc5023, NRHc5024, and NRHc5024A (all VNBI strains), were identified with truncated Sgf29 proteins as a result of either frameshift or point mutations resulting in premature stop codons (Fig. 5A). An additional three strains, NRHc5044 (VNI), PMHc1023 (VNII), and NRHc5030 (VNBI), were identified with missense mutations within the coding region of *SGF29* that were predicted to be deleterious by the online tools PROVEAN and PolyPhen-2 (Fig. 5B). In addition, tandem TA repeats ranging from 5 to 9 copies were identified in the second intron of *SGF29* (Fig. 5C; Supplementary Table S1). The number of repeats was significantly greater in VNBI, previously found to be the most genetically diverse and the most virulent genotype, compared to all other genotypes (p<0.001) (5).

**Figure 5:**
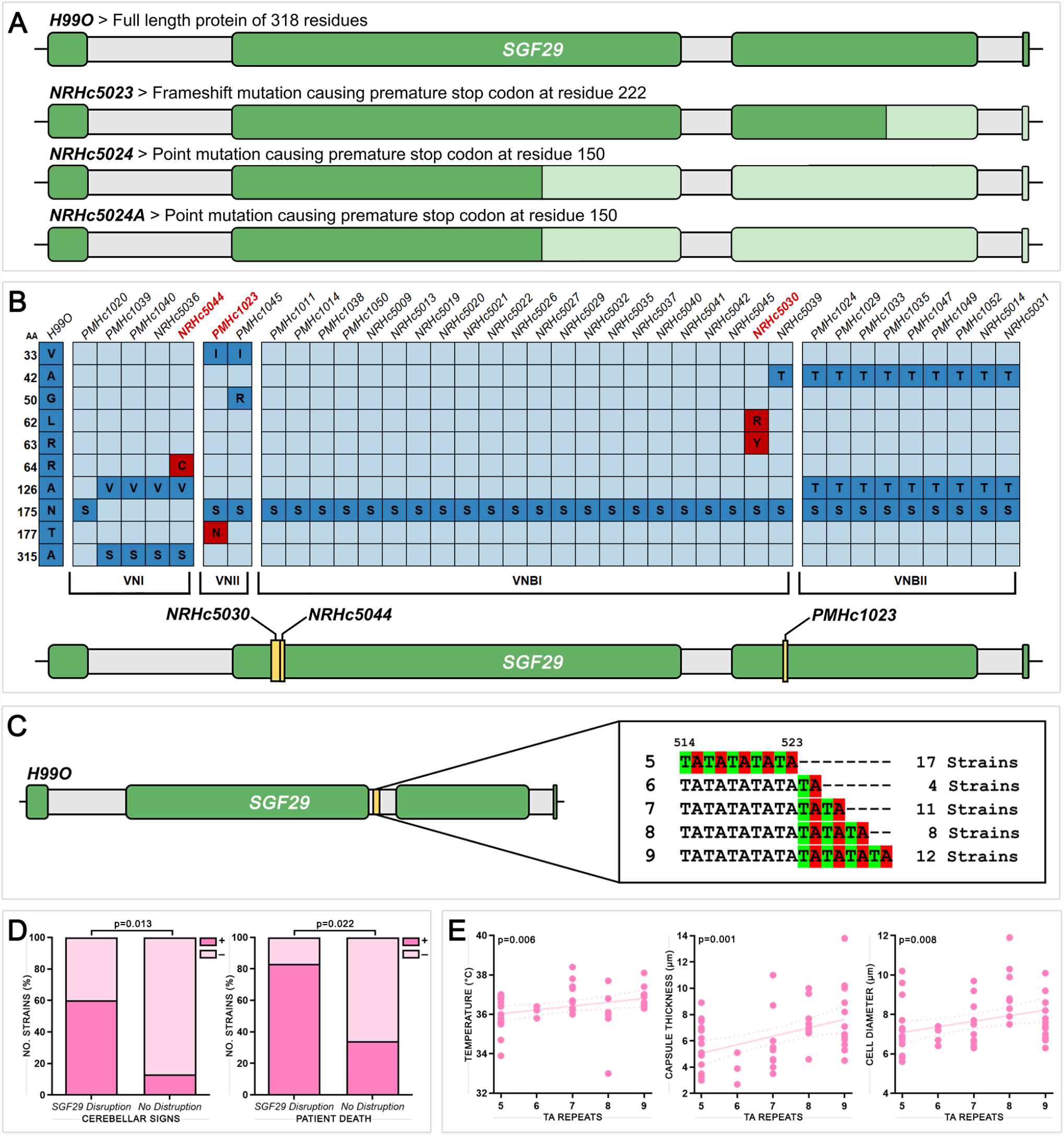
Mutations in *SGF29* in clinical isolates correlate with clinical and phenotypic variables. **(A)** Strains with mutations occurring in the *SGF29* gene resulting in truncated proteins. Lighter green sections indicate areas of deletion. **(B)** Strains with missense mutations occurring in the coding regions of the *SGF29* gene (top) and the location of the mutations (bottom). Dark blue boxes indicate mutations predicted to be neutral while red boxes indicate mutations predicted to be deleterious. **(C)** Tandem TA repeats ranging from 5 – 9 copies identified in the second intron of *SGF29*. **(D)** Significant correlations between clinical variables and strains possessing disruptions in *SGF29* (including both mutations resulting in truncated proteins and deleterious missense mutations). **(E)** Significant correlations between clinical and phenotypic variables and the number of TA repeats in the second intron of *SGF29*.

Correlations were assessed between either loss-of-function mutations or TA repeat number and the clinical and phenotypic variables recorded for these strains. Significant correlations were found between loss-of-function mutations, cerebellar signs (p=0.013), and patient death (p=0.022), indicating that Sgf29 disruption affects hypervirulence in clinical isolates (Fig. 5D). Additionally, of these six strains, five produced micro cells and shed capsule at high frequencies that were comparable to the hypervirulent H99 strains, while only one produced giant cells, and none produced irregular cells. For the TA repeat, significant correlations were found between more TA units and higher patient temperature (p=0.006), as well as greater capsule (p=0.001) and cell body (p=0.008) size (Fig. 5E).

## DISCUSSION

### The capacity to be pleomorphic aligns with virulence

Many studies have sought to correlate individual phenotypes with virulence, the most common being capsule size; however, these attempts have yielded substantially different results. For example, recent studies have reported larger capsules led to greater virulence in zebrafish (20), that smaller capsules resulted in increased macrophage uptake and decreased long-term patient survival (21), and that there was no association between capsule size and virulence in mice (22). Other studies have found strains with increased capsule shedding were associated with higher patient mortality (23), and that neurovirulence may be related to the total amount and speed of accumulation of capsule in the brain rather than the capsule size of individual cells (24). The disparate nature of these results likely reflects the fact that rather than a simple association with a single phenotype driving virulence, it is likely to be a combination of multiple phenotypes that determines the overall virulence profile of a strain.

In our previous study on Botswanan clinical isolates, we found significant correlations between phenotype and clinical variables suggesting a possible relationship between the two, including an association between the production of a range of cell variants and host death (5). However, the heterogenous nature of the dataset made determining causation difficult. In the current study looking at a set of strains with high genetic similarity, we have found the hypervirulent strains to be associated with larger average capsule, increased cell size variation, and increased production of micro cells, released capsule, and clustered capsule (Table 1). Taken together, the results of these two studies suggest that, rather than the production of any one phenotype, it is the capacity to be variable and produce a diverse range of phenotypes that is a potential driving factor behind increased virulence. In addition to the individual phenotypes themselves being associated with varying routes of pathogenesis, increased morphological heterogeneity as a whole may also contribute to virulence via immune escape.

Most of these variant phenotypes are still largely unexplored, with their presence in samples often not determined, or noted but not included in analyses. Micro cells, although having been reported as a common feature of cryptococcal infection (4, 17), and irregular cells, although observed in dozens of strains across *C. neoformans* and the *C. gattii* complex (5, 25), remain understudied. This may stem from the fact that most analyses of *Cryptococcus* are done using media that does not provide similar stresses to those encountered during mammalian infection and is therefore unable to induce these phenotypes. Similarly, while giant/titan cell types have become well-known in the cryptococcal community, a reliable way of inducing them *in vitro* was only recently developed (13).

### Microevolution and epigenetic plasticity enable phenotypic variation

It is well established that many fungal species including *Cryptococcus* can readily undergo morphological transitions in response to different environmental stimuli (26). What is less well understood are the exact mechanisms driving this morphogenesis and how it can affect virulence and the progression of disease. Microevolution is a term that typically refers to heritable changes accumulated over a relatively short period of time. The microevolution of cryptococcal strains over the course of infection has been well documented (27–30); an unsurprising fact given that *Cryptococcus* can establish chronic infections that can last months to years, during which the organism is subject to selection by host defence mechanisms (31). While this kind of microevolution often involves changes to the genome, instances of serial isolates displaying strong phenotypic changes with no or very limited genetic changes have also been noted (32). This suggests that epigenetic mechanisms may be involved that regulate gene expression and cellular response without altering the DNA sequence.

In the hypervirulent H99L lineage strains, the 734 bp deletion in *SGF29* is thought to play a role in their development of hypervirulence, presumably through the loss of histone H3K9 acetylation altering gene transcription across the genome. Studies looking at epigenetic regulation in *Cryptococcus* have found links between histone deacetylases and a range of cellular functions including thermotolerance, capsule formation, melanin synthesis, protease activity (33), and drug sensitivity (34), however there is still relatively little known about the role of epigenetics in regulating genome plasticity (35). In *Candida*, more direct links have been established with a recent study observing dynamic heterochromatin remodelling in response to changing the growth temperature from 30 °C to 39 °C, indicating that differential chromatin states controlling gene expression are linked to rapid adaptation (36).

In the H99OΔ mutant, we found substantially increased production of micro cells, released capsule, and clustered capsule, consistent with its switch to the hypervirulent phenotype (Table 1; Fig. 4) (9) and indicating that the loss of *SGF29* was indeed responsible for the enhanced variant production in the H99L lineage. However, this increased production of variants was retained in the complemented H99OΔ+ strain. Although unexpected, this result aligns with the previous finding where the reintroduction of *SGF29* was not sufficient to abolish hypervirulence in H99OΔ (9) and strongly indicates that epigenetic memory, which is difficult to restore, is involved in both the switch to hypervirulence and the production of variant phenotypes. Furthermore, removal and reintroduction of *SGF29* in an H99S background had no appreciable effect on either virulence (9) or morphological variation, and it appears that other mechanisms underly the increased virulence of this lineage. In contrast to the other hypervirulent strains which are through to have arisen during routine passaging, H99S acquired hypervirulence during passage through an animal model. This, coupled with the fact that H99S possesses no genetic mutations that can account for its hypervirulence, suggests that it has undergone substantial epigenetic change during its creation that is independent of the SAGA complex.

Also of interest was the observation that that hypervirulent strains H99S, H99F, KN99**a** and KN99α had increased susceptibility to azole drugs (Table 1) indicating a possible role of Sgf29 in the response to azole antifungals. Azole drugs function through inhibiting ergosterol synthesis, but while Sgf29 has been observed to alter the expression of ergosterol biosynthesis gene *ERG5* in some strains, it appears to have no effect in others (9). As an epigenetic regulator, the effects of the SAGA complex that includes Sgf29 can be indirect and wide-ranging and are therefore difficult to predict. Further study using ‘omics approaches would be very useful for understanding how these strains of variable virulence and phenotype respond to environmental challenges.

Given the apparent association between the loss of Sgf29, virulence, and the presence of morphological variants we extended our analysis to the clinical isolate set from our previous study on HIV/AIDS patients in Botswana (Fig. 5). Substantial protein truncations and deleterious mutations disrupting *SGF29* were identified in six of the 52 clinical strains (11.5%), and the presence of these loss-of-function mutations was found to be significantly correlated with patient death, suggesting hypervirulence. Additionally, a polymorphic TA repeat ranging from 5-9 copies was identified in the second intron of *SGF29* in the clinical isolates. Short tandem repeats are common in eukaryotes including fungi (37). While these mostly occur in noncoding regions and are not predicted to affect phenotype, the presence of self-complementary dinucleotides in introns has been shown to alter gene splicing, presumably through the formation of RNA hairpins, with splicing efficiency inversely correlated with repeat number (38). Here, increasing TA units were found to be significantly correlated with increased patient temperature and greater cell and capsule sizes, indicating a potential link with the regulation and expression of *SGF29*. These findings, together with the previous discovery of *SGF29* loss-of-function mutations in clinical isolates from the USA and India (9), provides further evidence for the importance of Sgf29 in a clinical setting.

### The morphological variants observed in this study have distinct cellular properties

Electron microscopy of the cryptococcal cell variant phenotypes, revealed new insights into their internal structure. The few giant cells observed here share similarities with titan cells reported in the literature, containing multiple small vesicles per cell and/or a single enlarged vacuole (12, 13). The giant cell shown in Supplementary Fig. 1A has a cell wall of 820 nm, which is larger than some reports of titan cells (314.7 ± 64.0 nm) (13) but smaller than others (2 – 3 μm) (17). These differences may be due to the inducing conditions and not inherent to the cell type itself. SBF-SEM revealed micro cells have an internal structure with organelles that seem similar to those seen in regularly sized cells and are distinct from shed capsule. This together with the fact that their size is outside the range of regular cell sizes, suggests that micro cells are a distinct cell type produced by regularly sized cells and are not simply at the small end of a spectrum of regular cell sizes.

Two types of capsule shedding were noted in this study: released capsule that was present in the medium and clustered capsule that appeared to be attached to otherwise regular cells (Fig. 3). There is currently little known about the capsular release mechanism of *Cryptococcus* and whether it is an active phenomenon or merely nonspecific shedding (39), although some studies indicate that capsular and exopolysaccharide materials originate from different pools (40). Some of the clustered capsule observed by light microscopy (Fig. 3A) appeared to have very small amounts of material contained within it that might be picked up inside the cell or pinched off as it exited the cell, implying an active phenomenon. However, in other cases capsule clusters appeared to have burst from the outer capsule, and it is quite possible that there are multiple capsule release mechanisms that are yet to be elucidated.

In our prior study of Botswanan clinical isolates we found irregular cells with elongated cell morphologies to be significantly more likely to occur in isolates obtained from patients who had undergone antifungal therapy prior to admission. Their presence was also negatively correlated with patient death, suggesting they may either be defective cells or a form of persister cell able to establish a chronic but non-lethal infection (5). Here, we observed them to have substantially thinner cell walls than even micro cells (Supplementary Fig. 1), which supports the idea that they may be more fragile and possibly defective. These cells share similar properties to a recently described group of cells termed titanides, characterised by a smaller cell size (2 – 4 μm), oval shape, and thinner cell walls. However, while titanides appear to be derived from titan cells, this is unlikely for the irregular cells observed here as they were present in samples that did not contain giant cells (13). Irregular cells were produced the most frequently by the hypervirulent congenic strain pair KN99**a** and KN99α, which displayed increased susceptibility to azole drugs, and also produced the highest levels of other morphological variants. This raises the question of whether this phenotype might be a consequence of the cell machinery producing large numbers of variants. More research into the exact nature of irregular cells will help us understand whether they are a phenotype of benefit to the cell or a defective by-product of phenotypic transition.

## Conclusion

Using a set of genetically similar but phenotypically diverse strains, we have shown a potential link between the capacity for phenotypic plasticity and virulence. We have provided further evidence that Sgf29 is an important epigenetic regulator that is likely involved in the transition to a hypervirulent phenotype, and have shown that loss-of-function mutations in *SGF29* may also be linked to virulence in clinical strains. Other pathways to increased virulence and pleomorphism also occur, however, as demonstrated with H99S. Together, our results demonstrate an impressive ability of *Cryptococcus* to produce variation even when growing as a clonal lineage, which may account for its capacity to produce a persistent, chronic infection in mammalian hosts.

## MATERIALS & METHODS

### Test strains

A collection of nine derivatives of *C. neoformans* strain H99 (8, 10) were used in this study, with details in Fig. 1 and Table 1. *SGF29* mutant strains described in (9) were also used, with details in Table 1. Botswanan clinical isolates described in (5) were used for bioinformatic analysis, with details in Supplementary Table S1.

### Growth curves and capsule induction

Culture conditions were as described in (5). Before experiments, cells were collected from overnight cultures by centrifugation and washed once with PBS. For growth curves, 10^3^ cells were inoculated into 2.5 ml of Sabouraud Dextrose Broth (SDB) in a 12-well tissue culture plate. Plates were incubated at 30 °C with 180 rpm shaking and cell density was measured every 2 hours by absorbance at 600 nm for a total of 48 hours. For capsule induction, 10^5^ cells were inoculated into 5 ml of Dulbecco’s Modified Eagle Medium (DMEM; Life Technologies) in a 6-well tissue culture plate and incubated with 5% CO_2_ at 37 °C for 5 days.

### Staining and microscopy

Staining, microscopy, and measurements of cell and capsule size were performed as described in (5) with slight modifications. Cells with *d*_*y*_ greater than 10 μm or less than 2 μm were identified as giant cells or micro cells, respectively. Giant cells, micro cells and irregular cells along with extracellular capsule were classified as morphological variants and were enumerated independently. For each individual strain, the relative frequency of micro cells, giant cells, released extracellular capsule, and clustered extracellular capsule was determined by looking over 20 random fields of view containing a minimum of 100 regular cells in total at 40x magnification and a category was assigned denoting either absence (**–**) or presence at one of three relative frequency levels (**+** / **+ +** / **+ + +**). Irregular cells were calculated as a % of the regular cell population including at least 100 cells.

### *Galleria mellonella* infection assays

*G. mellonella* infection assays were performed as described in (41). Briefly, for each test group 10 larvae were injected with an inoculum of 10^8^ cells, placed into a clean petri dish and incubated at 35 °C. The survival or death of each larva was recorded at 24-hour intervals over a period of 10 days.

### Antifungal susceptibility assays

Antifungal susceptibility assays were performed as described in (41) and in accordance with CLSI guideline M27-A3 for yeasts (42). Drugs were assayed at concentration ranges of 0.0039 – 4 μg/ml for AMB, NYS, ITC and VRC, and 0.0625 – 64 μg/ml for FLC and 5FC. Plates were incubated without agitation at 35 °C for 72 hours.

### Scanning electron microscopy

For standard scanning electron microscopy (SEM), sample preparation and visualisation were performed as described in (41). For serial block-face scanning electron microscopy (SBF-SEM), a single block was produced for analysis by mixing together aliquots from strains known to produce appreciable numbers of the different cell phenotypes (H99E, H99W, H99O, H99S, and KN99**a**) following capsule induction. Cells were harvested and washed with 0.1M phosphate buffer (PB), covered with 1 ml of fixative (2.5% glutaraldehyde, 2% PFA, 0.1 M PB) and left overnight at 4 °C. Secondary fixation, staining, ethanol dehydration, resin infiltration, and embedding were performed as described in (18). The final sample block contained approximately 5000 cells. The sample block was then loaded onto a Zeiss Sigma Variable Pressure SEM equipped with Gatan 3View 2XP. Inverted backscattered electron images (8192 × 8192 pixels, 16-bit) were acquired with XY pixel size of 100 nm, and Z pixel size (slice thickness) of 10 nm.

### Bioinformatic analysis of clinical isolates

Sequencing data were downloaded from the NCBI Sequence Read Archive and assembled and analysed using Geneious 6.0.6. The effect of missense mutations on the functionality of the protein was predicted using the PROVEAN (43) and PolyPhen-2 (44) online tools and validated with assistance from the PRALINE tool (45) to confirm changes to hydrophobicity.

### Statistical analysis

Significant differences between treatment groups for *Galleria* infection assays were determined using the log-rank test to compare the distributions between each pair of strains. Significant differences between strains for phenotypic data were determined using two-tailed unpaired t tests with Welch’s correction and differences in variance were assessed by F test analysis. For correlations between genetic, clinical, and phenotypic data, tests between continuous variables used Spearman rank-order correlations, those between continuous and binary variables used Mann-Whitney U tests, and those between binary variables used chi-square tests, or Fisher’s exact tests if any expected value was <5. P values of <0.05 were considered significant. Error bars represent the mean +/- 95% confidence intervals. Data were analysed using Excel (Microsoft Corporation), Prism 5 (GraphPad Inc.), and SPSS Statistics (IBM) software.

## ACKNOWLEDGEMENTS

The authors acknowledge the facilities as well as the scientific and technical assistance of the Microscopy Australia (micro.org.au) node at the University of Sydney: Sydney Microscopy & Microanalysis. We thank Christina Cuomo and Poppy Sephton-Clark (Broad Institute) for their assistance with genomic data analysis.

## AUTHOR CONTRIBUTIONS

KEF and DAC conceived and designed the experiments. KEF produced, collated, and analysed the data and wrote the manuscript with assistance from DAC and JAF.

## Figure Legends

**Supplementary Figure 1: Morphological variants produced by H99 strains visualised by electron microscopy**. Serial block face scanning electron microscopy (SBF-SEM) images of a representative **(A)** giant cell, **(B)** regular cell, **(C)** micro cell, and **(D)** irregular cell present in H99 derivative strains. White arrows indicate location of cell wall, black arrow indicates location of vesicle cluster. **(E)** Scanning electron microscopy (SEM) images showing regular cells grown under standard conditions (left) compared to cells grown under inducing conditions with large quantities of clustered capsule (right). **(F)** SEM images showing that a bleb of shed capsule (left) is clearly distinct from a micro cell (right). **(G)** Micro cells being released from regular cells; SEM image of two approximately 1.5 μm micro cells emerging from a regularly sized KN99**a** cell covered in burst capsule (left), and SBF-SEM image of an approximately 1 μm micro cell seen budding from a regularly sized cell (right).

## Table Headings

**Supplementary Table S1:** Genetic, clinical and morphological data for the Cryptococcus neoformans strain collection from Botswana

